# Analysis of FAIMS for the Study of Affinity-Purified Protein Complexes Using the Orbitrap Ascend Tribrid Mass Spectrometer

**DOI:** 10.1101/2024.12.02.626431

**Authors:** Pratik Goswami, Joseph Cesare, Michaella J. Rekowski, Zachary Clark, Janet Thornton, Michael P. Washburn

**Author notes:** To whom correspondence should be addressed: Michael P. Washburn, Ph.D., Department of Cancer Biology, University of Kansas Medical Center, Kansas City, KS, 66160, USA.

## Abstract

In this study, we analyzed the combination of affinity purification mass spectrometry (AP-MS) with high-field asymmetric waveform ion mobility spectrometry (FAIMS), integrated between nanoLC-MS and an Orbitrap Ascend Tribrid Mass Spectrometer. Our primary objective was to evaluate the application of the FAIMS interface for detecting affinity purified SAP25 protein complexes with enhanced sensitivity and robustness. As a result, we observed that nanoLC-FAIMS-MS (with FAIMS) significantly improved the sensitivity and detection limits at the protein level, peptide level and significantly reduced chemical contaminants compared to nanoLC-MS alone without FAIMS (No FAIMS). This FAIMS configuration resulted in 42% and 92% increases for the total proteins and unique proteins, respectively, and 44% and 88% increases for total peptides and unique peptides compared to the No FAIMS configuration. Our in-depth comparison of FAIMS and No FAIMS shows that FAIMS outperforms by significantly reducing the missing value by <15% in datasets and plays a significant role in filtering chemical contaminants. Our findings highlight the potential of FAIMS with Orbitrap Ascend Tribrid Mass Spectrometer to enhance the depth of AP-MS analysis. The data were deposited with the MASSIVE repository with the identifier MSV000096548.

## Introduction

Comprehensive proteomic profiling, which involves characterizing thousands of proteins, remains a significant challenge ^1^. Despite technological advancements, the complexity of the proteome, a dynamic range of protein expression, issues with detecting low-abundance proteins, and the complexity of data have persisted as major obstacles ^2^ ^3^ ^4^. Liquid chromatography coupled with mass spectrometry (LC-MS) is a key technique used to address these challenges, providing in-depth proteome profiling across various sample types including tissues, exosomes, and cells ^5, 6^. LC-MS is widely utilized for its effectiveness in managing proteome complexities, making it essential for detailed profiling of protein levels, functions, interactions, and modifications. ^7, 8^. One of the key applications of LC-MS is in affinity purification mass spectrometry (AP-MS), which facilitates the exploration of protein-protein interactions. In AP-MS, affinity-tagged bait proteins are isolated along with their interacting “prey” proteins, followed by mass spectrometric analysis to elucidate these interactions ^9–11^. Studying these interactions is essential for understanding cellular processes and disease mechanisms, making AP-MS a powerful tool in basic and applied research ^10, 12^. Advances in mass spectrometry, such as high-resolution analyzers, have enhanced the accuracy and sensitivity of AP-MS, enabling deeper insights into complex biological systems ^8^.

Sin3 associated protein, SAP25 is a member of the Sin3/HDAC complex, playing a vital role in transcriptional repression. Previous studies suggested that SAP25 acts as an adaptor protein and is associated with O-linked N-acetylglucosamine transferase (OGT) ^13^, SCF E3 ubiquitin ligase complex SKP1/FBXO3/CUL1 and the ubiquitin carboxyl-terminal hydrolase 11 (USP11). These proteins play a vital role in transcription regulation, by changing post-translational modifications and DNA alterations. Notably, these enzymatic and regulatory complex proteins are uniquely associated with SAP25, among the other eleven Sin3 Complex proteins ^14^. This makes SAP25 important to study by AP-MS. When studying protein complexes by AP-MS it is important to take into consideration the possible protein contaminants that can occur from using specific affinity purification methods ^15^. In addition it is also important to take into consideration the chemical contaminants that can interfere with peptide analysis by mass spectrometry ^16^.

Nanoscale capillary liquid chromatography with mass spectrometry (nanoLC−MS) remains a cornerstone in mass spectrometry-based proteomics ^17^. Despite its central role, nanoLC−MS faces limitations, particularly in requiring larger sample volumes, is time-consuming, and becomes increasingly challenging when analyzing many samples ^18^. High-field asymmetric waveform ion mobility spectrometry (FAIMS) enhances proteomics by reducing background interference and minimizing contaminants, thus improving reliability and sensitivity ^18, 19^.

FAIMS works at atmospheric pressure and separates ions based on their various mobility in low and high electric fields ^20^. Ions travel between two parallel electrodes, where an asymmetric waveform is applied, causing displacement due to their differential mobility. This displacement is corrected by applying the compensation voltage, CV ^21^. Additionally, FAIMS offers gas-phase fractionation through multiple compensation voltages (CVs), which effectively deconvolutes overlapping peptide signals, providing significant advantages over traditional nanoLC−MS approaches ^1^. When combined with mass spectrometry, the FAIMS separates ions before they enter the vacuum chamber of the mass spectrometer, reducing chemical background interference and enhancing signal-to-noise ratios. ^22^. Previous studies have highlighted the benefits of FAIMS in proteomics applications, including top-down studies and single-shot proteomics. ^1, 18, 23–27^. However, there is limited comparative research on FAIMS for bottom-up proteomics with affinity-purified protein complexes^28^.

Technological developments in mass spectrometry (MS)-based proteomics continually enhances the detection capability and precision of proteome analyses ^29^. The LTQ Orbitrap, introduced by Thermo Fisher Scientific in 2005, marked a significant milestone. Over the past decade, subsequent advancements have focused on enhancing sensitivity and scan speed, addressing the evolving demands of proteomics ^27^. However, achieving comprehensive proteome coverage, sensitivity and speed are still major challenges ^30, 31^. The Orbitrap Ascend, equipped with a linear ion trap, quadrupole, and Orbitrap, addresses these challenges by improving scan speed and sensitivity ^31, 32^. Recent comparative studies by Shuken et al. and He et al. have demonstrated that the Orbitrap Ascend outperforms the Orbitrap Eclipse in both protein detection and quantification, further underscoring the potential of these technologies in proteomic research ^30, 31^.

In this study, we present the application of an advanced FAIMS device integrated between nanoLC-MS on the Orbitrap Ascend Tribrid Mass Spectrometer for a bottom-up proteomics approach targeting affinity-purified protein complexes. First, we carried out SAP25 affinity purification from Flp-In™-293 cells stably expressing Halo-SAP25. The purified proteins were then subjected to digestion into peptides followed by mass spectrometry with FAIMS and No FAIMS configuration. (Figure 1) ^33^. In this study, we successfully characterized Halo-SAP25 purified proteins with both FAIMS and No FAIMS configurations. Our findings revealed that FAIMS enhances the detection of a number of proteins and peptides. Next, we tested FAIMS efficacy to filter the potential contaminants and reduce the missing values. Overall, our results demonstrate that the distinct effectiveness of FAIMS substantially improves the detection of affinity-purified protein complexes, especially the lower abundance proteins and peptides, while significantly reducing the level of contaminants and the percentage of missing values when used with the nanoLC on Orbitrap Ascend Tribrid Mass Spectrometer.

**Figure 1.**
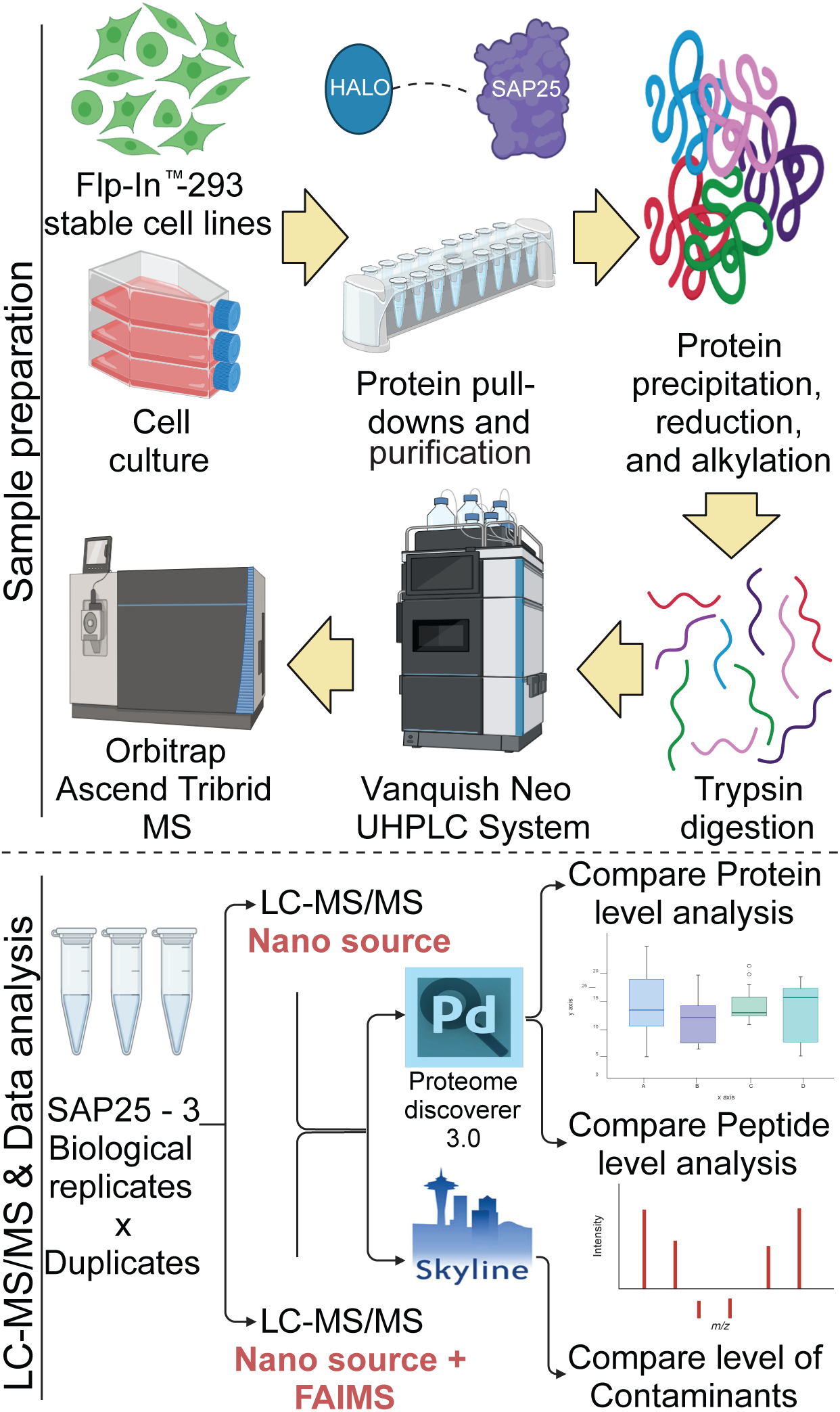
Experimental workflows for sample preparation in LC-MS/MS quantitative proteomics strategies. Cell extracts were prepared from Flp-In™-293 stable cell lines expressing Halo-SAP25 protein. After lysis, proteins were separated using Halo-tag magnetic beads. The resulting elute was subjected to protein precipitation, reduction, alkylation, and trypsin digestion. The peptides were subsequently injected using the Vanquish Neo (Thermo) nano-UPLC followed by mass spectrometry. The detection was compared by nanoLC-MS and nanoLC-FAIMS-MS to determine the benefits of FAIMS for the proteins, peptides, and contaminants detection. The raw files for the proteins and peptides were analyzed using Proteome Discoverer 3.0 and contaminants were analyzed by Skyline 24.1 Finally, statistical analysis was conducted to identify differences in detection between FAIMS and No FAIMS configurations.

## Materials and Methods

Magne® HaloTag® beads (G7281), AcTEV protease (#12575015), Pierce TCEP-HCI (#20490), DSSO (disuccinimidyl sulfoxide) (#A33545), and Formic Acid, LC-MS Grade (#28905) were purchased from Thermo Fisher Scientific. The reagents 2-Chloroacetamide (#C0267), Triton X-100 (#T8787), Urea (#U1250-1KG), and Trichloroacetic acid solution (#T0699) were obtained from Sigma. rLys-C, Mass Spec Grade (#V1671), protease inhibitor cocktail (#G652A) and Sequencing Grade Modified Trypsin (#V5117) were sourced from Promega. Salt Active Nuclease High Quality (#70920-202) was acquired from ArcticZymes.

### Preparation of whole cell lysates

A Flp-In™-293 cell line stably expressing Halo-SAP25 regulated by a CMV promoter was established as described previously (Banks et al., 2018). Approximately 1.5-6.3 × 10^8 cells were harvested, washed twice with ice-cold PBS and lysed by using Dounce homogenization in ice-cold lysis buffer containing 20 mM HEPES (pH 7.5), 1.5 mM MgCl2, 0.42 M NaCl, 10 mM KCl, 0.2% Triton X-100, and 0.5 mM DTT, 1X protease inhibitor cocktail (G652A) and 500 units of salt-active nuclease (SAN). The lysates were incubated for 2 hours at 4°C. After incubation, the lysates were centrifuged at 40,000 x g at 4°C for 30 minutes. The salt concentration of the resulting supernatants was reduced to 0.3 M NaCl by adding ice-cold buffer containing 10 mM HEPES (pH 7.5), 1.5 mM MgCl2, 10 mM KCl, 0.5 mM DTT, and 1X protease inhibitor. The lysates were centrifuged again at 40,000 x g at 4°C for 30 minutes. The resulting supernatant was collected for Halo affinity purification and subsequent cross-linking experiments.

### Purification and preparation of recombinant protein complexes

For Halo-Tag purification, lysates were incubated overnight at 4°C with Magne® HaloTag® beads prepared from 0.2 ml of bead slurry. The beads were isolated using a DynaMag-2 Magnet and the supernatant was discarded. Beads were washed 4 times in wash buffer containing 10mM HEPES (pH 7.5), 300mM NaCl, 10mM KCl, 1.5mM MgCl2, and 0.2% Triton X-100. Bound proteins were eluted from beads by incubating the beads for >2 hours at 4°C with 200 µl elution buffer having 50 mM HEPES (pH 7.5), 0.5 mM EDTA, 1 mM DTT, and 30 units AcTEV. The resulting eluate was collected, and proteins crosslinked in 5 mM DSSO for 40 minutes at room temperature. The crosslinking reaction was quenched with 50 mM NH4HCO3 for 15 minutes at room temperature.

### Digestion of proteins for mass spectrometry

Halo-purified proteins were incubated overnight at 4°C with 100% (w/v) ice-cold trichloroacetic acid (TCA) to a final concentration of 20%. The precipitated protein pellets were collected by centrifugation at 21,000 x g for 30 minutes at 4°C. The pellet was washed two times with acetone, air dried, and resuspended in 8M urea dissolved in 100 mM Tris-HCl (pH 8.5). The formation of Disulfide bonds was reduced and alkylated with Tris(2-carboxyethyl)-phosphine hydrochloride (TCEP) and chloroacetamide (CAM) subsequently. Denatured proteins were initially treated with 0.1 μg of Lys-C for < 6 hours at 37°C. The urea concentration was lowered to 2M by adding 100mM Tris-HCl, pH 8.5 followed by 2mM final concentration of CaCl2. At the end, proteins were cleaved with 0.5 μg of trypsin. The reactions were terminated by the addition of formic acid to a final concentration of 5% before proceeding with mass spectrometry analysis.

### Mass Spectrometry

Proteomic profiling of Halo-SAP25 isolated complexes was performed using a Vanquish Neo UHPLC system coupled with an Orbitrap Ascend Tribrid Mass Spectrometer (Thermo) for nanoLC-MS and nanoLC-FAIMS-MS analyses. Tryptic digests of SAP25 samples were introduced onto the C18 trap column (PepMap™ Neo Trap, 0.3 mm × 5 mm, 5 µm particle size) using the Vanquish Neo (Thermo) nano-UPLC. Before being injected into the mass spectrometer, peptides were moved onto the separation column (PepMap™ Neo, 75 µm × 150 mm, 2 µm C18 particle size, Thermo).

All the peptides were washed and loaded with a flow rate of 350 µL/min with 2% mobile phase B (mobile phase A: 0.1% formic acid in water; mobile phase B: 80% acetonitrile, 0.1% formic acid). Peptide separation occurred beyond 100 minutes from 2-25% mobile phase B. This was followed by an increase to 40% B over the next 20 minutes. Before the next injection with 2% B, the column was cleaned and equalized for 15 min with 100% B.

The nano-LC was directly connected with the Thermo Orbitrap Ascend Tribrid Mass Spectrometer. A silica emitter (20 µm i.d., 10 cm) integrated with a high field asymmetric ion mobility source (FAIMS). The data were acquired by data-dependent acquisition (DDA), and the intact peptides were detected in the Orbitrap with a resolving power of 120,000, covering the m/z range from 375 to 1500. Higher-energy collisional dissociation (HCD) at 28% normalized collision energy (NCE) was employed to fragment peptides with charge states ranging from +2 to +7 with dynamic exclusion set for 60 seconds after the initial detection of a precursor ion.

In terms of FAIMS, the compensation voltage (CV) were selected as -45V (1.4 s), -60V (1 s), and -75V (0.6 s) for a time of 3s with source ionization +1700 V, with 305°C temperature of the transfer tube. The resulting raw data were searched in Proteome Discoverer 3.0 (PD; version 3.0; Thermo Fisher Scientific) against the human protein database of Uniprot 05-05-2023 with a common contaminants database. Protein and peptides abundances were exported to Microsoft Excel and graphs were prepared using GraphPad Prism.

### Experimental Approach and Data Analysis

The comparative analysis to identify Halo-SAP25 affinity-purified protein complexes with FAIMS and No FAIMS configurations required a minimum three biological replicates (Table S1). All three biological replicates were run twice, resulting in a total of six runs for each configuration. All the raw files were searched using Thermo Scientific Proteome Discoverer 3.0 software against the human protein database downloaded from Uniprot 05-05-2023, along with the contaminants database downloaded from Protein Center database service by Thermo Fisher Scientific. Trypsin was specified as the proteolytic enzyme, permitting up to two missed cleavages. For protein identification, the following parameters were used: Precursor mass error tolerance of 10 ppm and fragment mass error at 0.02 Da. Precursor masses with a minimum of 350 Da and a maximum of 5000 Da were permitted. Targeted FDR relaxed at 0.05 and strict at 0.01. Protein abundance values, Normalized and grouped Abundance of proteins along with their Observed Peptides and PSM values were exported as Excel file directly from the proteome discoverer. For comparative analysis of proteins observed under nanoLC-MS and nanoLC-FAIMS-MS, study factors were created in Proteome Discoverer. For Contaminants and grouped abundances count (no value) less than 1 were filtered out. Data were sorted out in Microsoft Excel and graphs were prepared using GraphPad Prism 9. Statistical analysis was performed using Welch’s t-test.

The comparative analysis to identify the level of contamination between Halo-SAP25 affinity-purified protein complexes with FAIMS and No FAIMS conditions was performed by importing the raw files to the Molecular Contaminants MS1 Filtering v4.1 document with the document’s default settings in Skyline-daily (64-bit) 24.1.1.284 (bc93c2813) ^34^ ^35^ ^36^. Synchronized peak integration based on peak area was performed between replicates with wide windows of integration to account for large elution profiles and the lack of molecular identity validation since these precursors were not selected for fragmentation. Skyline ^34^ ^35^ ^36^ results were exported and formatted with R Studio for Microsoft Excel.

## Results

### FAIMS Technology Enhances Protein Level Detection in Affinity-Purified Protein Sample

The primary aim of this study was to assess the impact of FAIMS in protein-level, peptide-level, and contaminants-level detection when used with nanoLC-MS. To evaluate the importance of FAIMS in protein detection, we initially developed a stable cell line expressing a recombinant version of SAP25 (NP_001335606.1) with an N-terminal Halo-Tag (Halo-SAP25) in Flp-In™-293 cells. The cells were lysed, followed by Halo affinity purification and trypsin digestion, as described previously ^14^. The isolated proteins were then injected into Vanquish Neo (Thermo) nano-UPLC directly interfaced with an Orbitrap Ascend Tribrid Mass Spectrometer. To compare the impact of FAIMS, we used Halo-SAP25 affinity-purified protein complexes with FAIMS and with No FAIMS configuration. Using a 140-minute instrument method, we performed six replicates for both No FAIMS and FAIMS at three compensation voltages (-45V, -60V, and - 75V), to assess their impact on protein fragmentation and detection (Table S1). In this study, we sought to investigate the advantages of the FAIMS interface for enhancing the identification and quantification of proteins, peptides, and contaminants (Figure 1).

The Venn diagram (Figure 2A) illustrates the overlap and uniqueness of proteins identified under both configurations. In the FAIMS-assisted analysis, a total of 385 proteins were identified, with 176 proteins uniquely detected under FAIMS configuration. In contrast, the No FAIMS configuration yielded the identification of 222 proteins, of which only 13 were unique to this setting. Notably, the FAIMS configuration resulted in a 13-fold increase in the number of unique proteins detected compared to the No FAIMS configuration. A total of 209 proteins were commonly identified under both FAIMS and No FAIMS configurations. To further assess the reproducibility and distribution between the SAP25 runs, we performed a Principal Component Analysis (PCA) on the proteomic data for both FAIMS and No FAIMS configurations. The PCA plot (Figure 2B) illustrates the distribution of SAP25 runs analyzed with FAIMS and with No FAIMS configuration. The first two principal components (PC1 and PC2), accounting for 53.9% and 19.0% of the variance respectively, clearly separate the samples processed with FAIMS from those processed with No FAIMS. The distinct and tight clustering of FAIMS and No FAIMS samples suggests a differential separation between the two methods. Both the Venn diagram and PCA plot were prepared using Proteome Discoverer (PD; version 3.0; Thermo Fisher Scientific). Additionally, when comparing the total number and abundance of proteins, we observed that FAIMS detected twice as many proteins, with a particular increase in low-abundance proteins (Table S1). Figure 2C illustrates the distribution of the average number of proteins across six runs for both FAIMS and No FAIMS configurations, with a total of 340 proteins detected with FAIMS (depicted in blue) and 158 proteins detected with No FAIMS (depicted in red). The data demonstrate a statistically significant increase in protein detection and normalized abundances when FAIMS is applied (Figure 2D). The normalized protein abundances (log2) for individual Halo-SAP25 affinity-purified protein complex runs, as shown in Figure 2E, indicates the reduction in normalized protein abundances (log2) with FAIMS configuration due to detection of lower abundance proteins (Table S1). These results underscore the significant enhancement in proteomic depth afforded by the incorporation of FAIMS.

**Figure 2.**
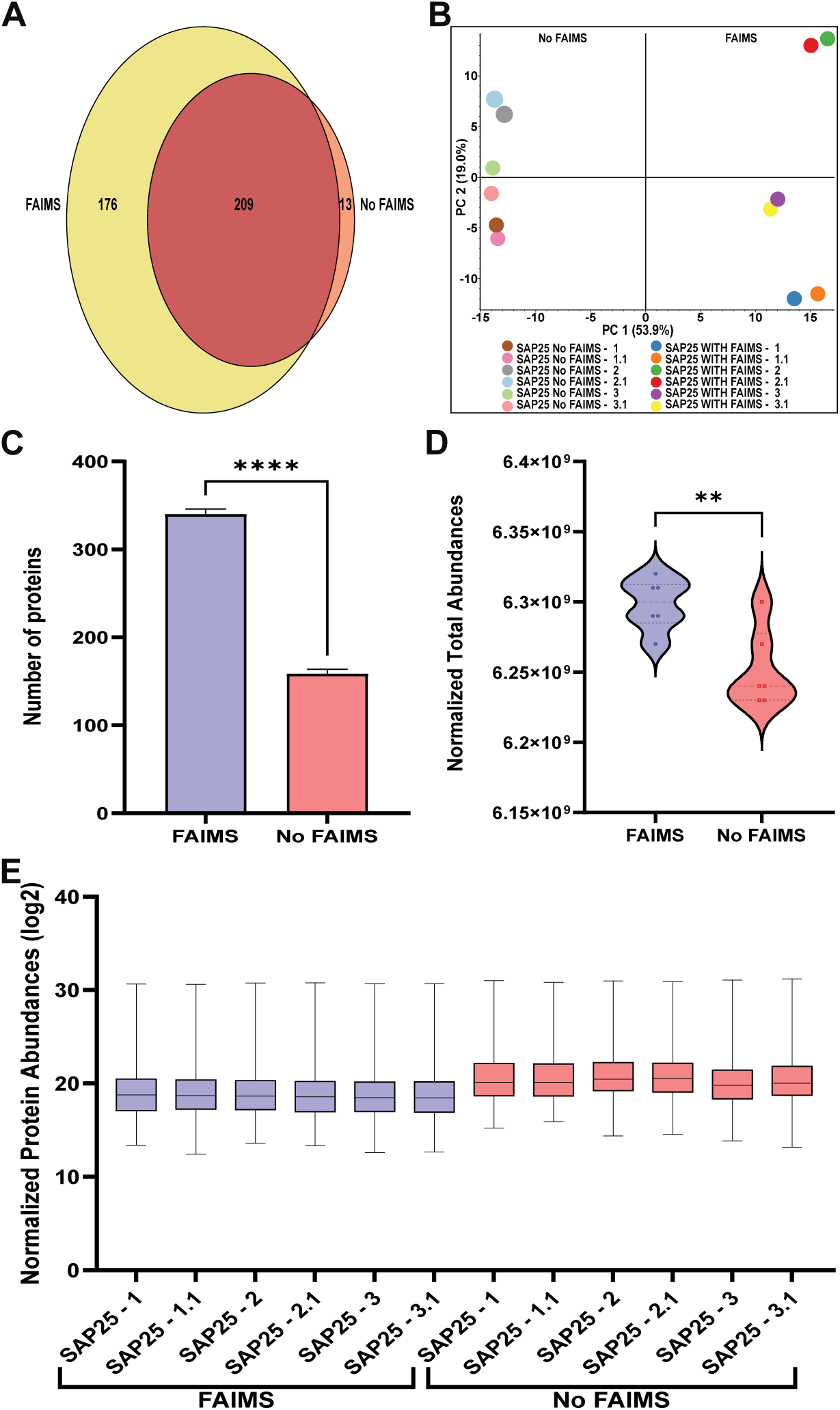
Comparative Analysis of Protein level Detection using FAIMS and No FAIMS Configurations. (A) Venn diagram illustrating the overlap in a number of SAP25 purified proteins observed with FAIMS and No FAIMS configurations. (B) Principal Component Analysis (PCA) plot showing the variance in protein detection across SAP25 runs under No FAIMS (colored circles on the left) and FAIMS (colored circles on the right) configurations. (C) Bar graph representing the total number of SAP25 purified proteins detected with FAIMS (blue) and No FAIMS (red) configurations. Values represent the averages of six runs, and Error bars indicate standard deviation. (D) Violin plot comparing the normalized total abundances of SAP25 purified proteins observed with FAIMS and No FAIMS configurations. Values represent the averages of six runs, and Error bars indicate standard deviation. (E) Box and Whisker plot representing log2 value of normalized SAP25 purified protein abundances observed across a total of twelve SAP25 runs under FAIMS and No FAIMS configurations.

### FAIMS Technology Enhances Peptide Level Detection in Affinity-Purified Protein Sample

In addition to enhancing protein level identification, we also focused on the impact of FAIMS on peptide-level detection (Table S2). The Venn diagram (Figure 3A) indicates that in the FAIMS-assisted analysis, a total of 1321 peptides were identified, with 761 peptides uniquely detected. In contrast, the No FAIMS configuration identified 729 total peptides, of which only 169 were unique to this setting, while 560 peptides were common with both FAIMS and No FAIMS configurations. The broader detection of unique peptides with FAIMS, as evidenced by the Venn diagram, is further corroborated by the PCA analysis (Figure 3B), which offers insights into the reproducibility and consistency of peptide identification across runs. These trends also align with those observed at the protein level. Both the Venn diagram and PCA plot were prepared using Proteome Discoverer (PD; version 3.0; Thermo Fisher Scientific). Quantitative comparison of peptide counts across six runs, as shown in Figure 3C, reveals that an average of 690 peptides were observed with FAIMS, while No FAIMS accounted for only 389 peptides. The use of FAIMS nearly doubled the number of peptides identified per sample, reflecting its significant contribution to the depth of proteome coverage. The data demonstrate a statistically significant increase in peptide detection and normalized total abundances when FAIMS is applied (Figure 3D). We observed a similar pattern in the individual normalized peptide abundances (log2) as seen with protein abundances, suggesting that FAIMS significantly enhances the detection of lower-abundance peptides (Figure 3E).

**Figure 3.**
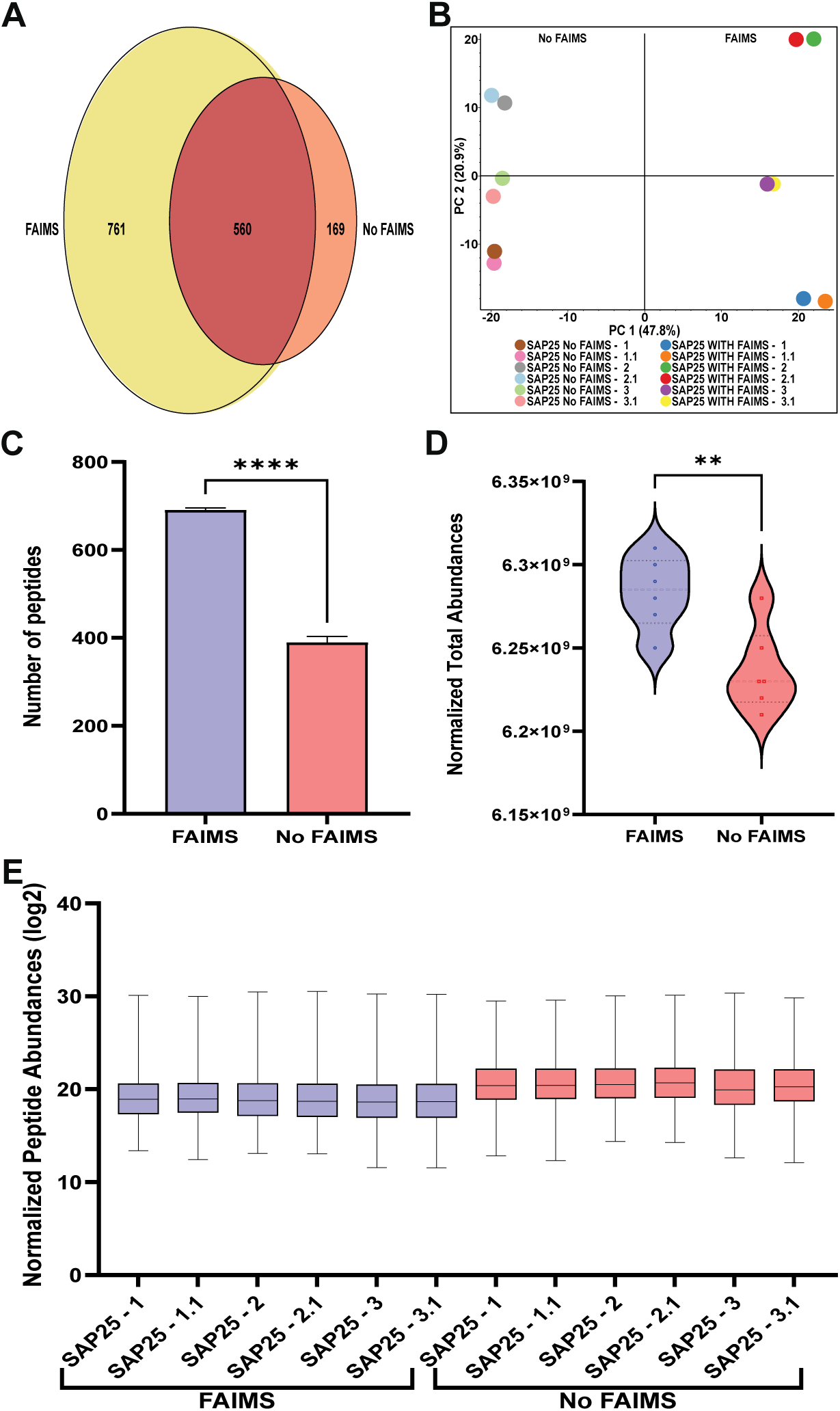
Comparative Analysis of Peptide level Detection using FAIMS and No FAIMS Configurations. (A) Venn diagram illustrating the overlap in a number of SAP25 purified peptides observed with FAIMS and No FAIMS configurations. (B) Principal Component Analysis (PCA) plot showing the variance in peptide detection across SAP25 runs under No FAIMS (colored circles on the left) and FAIMS (colored circles on the right) configurations. (C) Bar graph representing the total number of peptides detected with SAP25 purification with FAIMS (blue) and No FAIMS (red) configurations. Values represent the averages of six runs Error bars indicate standard deviation. (D) Violin plot comparing the normalized total abundances of peptide observed with SAP25 purification with FAIMS and No FAIMS configurations. Values represent the averages of six runs. Error bars indicate standard deviation. (E) Box and Whisker plot representing log2 value of normalized peptide abundances observed with SAP25 purification across multiple SAP25 runs under FAIMS and No FAIMS configurations.

Additionally, we have also considered unique and razor peptides for quantification to check the efficiency of FAIMS. Proteins identified with at least one unique peptide were considered. Our result showed that the sum of unique peptides observed with FAIMS is higher than observed with No FAIMS, which is 949 and 745 respectively. However, in both configurations, the majority of proteins were identified by a single unique peptide while the razor peptide count was 69 for FAIMS and 51 for No FAIMS (Figure S1 and Table S4).

### FAIMS Technology Significantly Reduces Missing values and identifies low-abundance proteins in Affinity-Purified Protein Sample

Missing values in proteomics data generally disrupt the integrity of statistical data analysis ^37 38 39^. In our study, we aimed to highlight the missing values detected in protein and peptide level among Halo-SAP25 affinity-purified protein complex runs between the FAIMS and No FAIMS configurations. To examine whether FAIMS plays a significant role in reducing the number of missing values, we performed hierarchical clustering with normalized log2 abundance, using One Minus Pearson Correlation as the metric and average linkage as the method. Our analysis revealed that a high number of SAP25 purified proteins with normalized log2 abundance > 0 were detected in the FAIMS groups compared to the No FAIMS group, where a greater number of missing protein values were observed (Figure 4A and Table S3).

**Figure 4.**
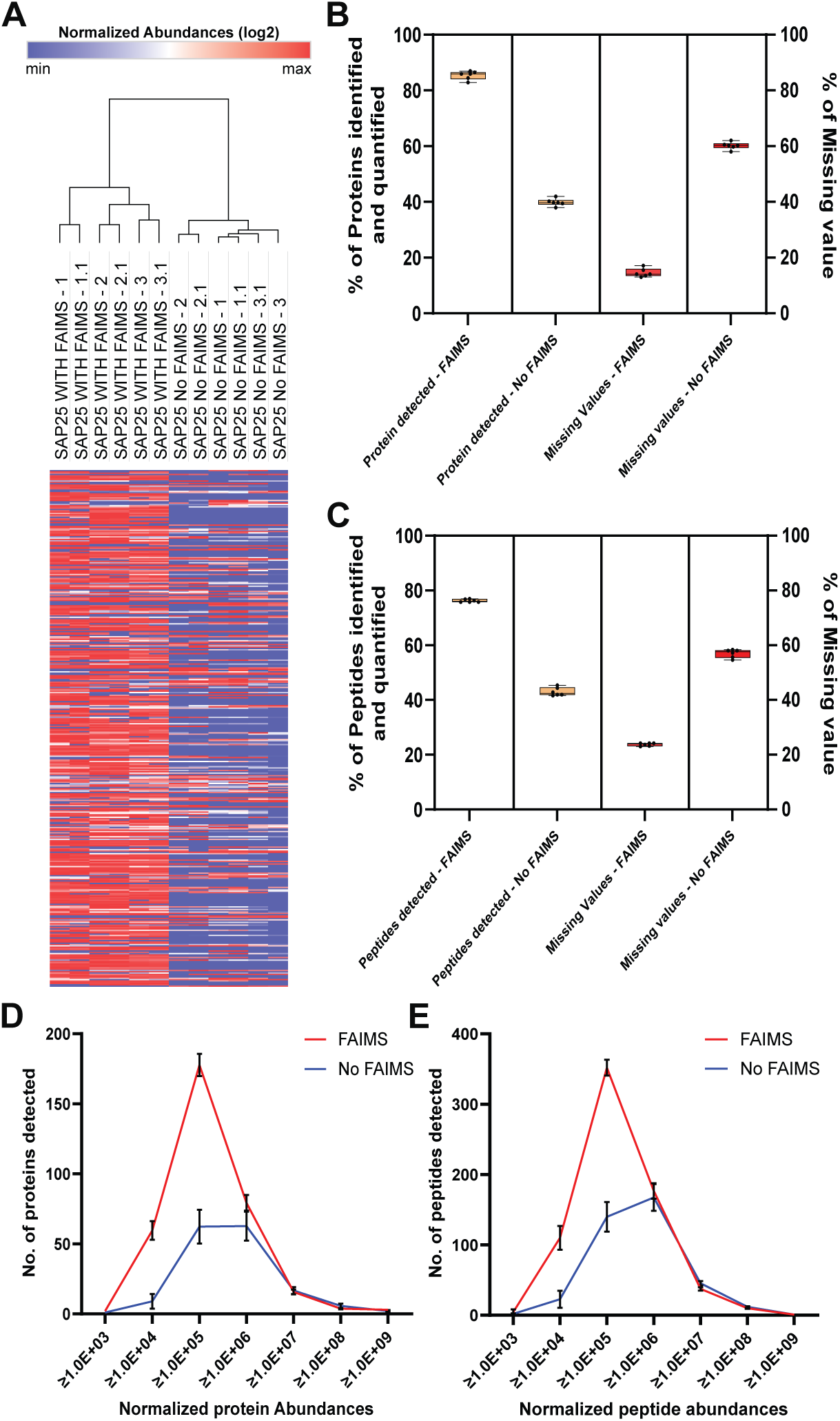
Association of FAIMS reduces missing values and improves data completeness. (A) Hierarchical clustering of Proteins purified from Flp-In™-293 cells expressing Halo-SAP25. The comparative heatmap was created using the Morpheus online tool (https://software.broadinstitute.org/morpheus). Values were log2 Normalized protein abundances with missing values set to zero. Clustering was performed with One minus Pearson correlation and average linkage. (B) Nested Box and Whiskers plot comparing the percentage of SAP25 purified proteins identified and quantified, as well as the percentage of proteins with missing values observed with FAIMS and No FAIMS configurations. (C) Nested Box and Whiskers plot comparing the percentage of peptides identified and quantified, and the percentage of peptides with missing values observed with SAP25 purification with FAIMS and No FAIMS configurations. (D) Comparison of protein detection with FAIMS (Red) and No FAIMS (Blue) configurations across various normalized abundance ranges. (E) Comparison of peptide detection with FAIMS (Red) and No FAIMS (Blue) configurations across various normalized abundance ranges.

To further confirm the differences in missing values between the two groups, we calculated the percentage of missing values observed with FAIMS and No FAIMS configurations at the protein and peptide levels (Table S1-S2). This was validated by analyzing the number of proteins identified and quantified, as well as those lacking abundance values in each configuration. In the FAIMS group, we found that on average, 85.43% of proteins were identified and quantified with only 14.57% of proteins showing missing values among the six runs of SAP25. In contrast, in the No FAIMS group, only 39.42% of proteins were identified and quantified while the remaining 60.58% of proteins had missing values (Figure 4B). A similar comparison was carried out with peptides, demonstrating the same pattern. In the FAIMS group, 76.25% of peptides were identified and quantified, with only 23.75% showing missing values. Conversely, in the No FAIMS group, 43.01% of peptides were identified and quantified, while the remaining 56.99% had missing values (Figure 4C). This observation further confirms the previous claims that FAIMS helps to minimize the number of missing values ^6, 40^. Next, we sought to determine how FAIMS significantly enhances the detection of lower abundance proteins and peptides. To verify that, we selected several abundance ranges, increased 10-fold, and analyzed the number of proteins and peptides detected within each range. Our result suggests that FAIMS substantially improves the detection of lower abundance proteins (Figure 4D) and peptides (Figure 4E), particularly with ranges ≥ 1.0E+03, ≥ 1.0E+04, and ≥1.0E+05, while at higher abundance ranges, there were minimal differences in numbers observed in FAIMS and No FAIMS configuration.

Among six Halo-SAP25 affinity-purified protein complex runs on average 2, 60, and 178 proteins and 4, 110, and 352 peptides were detected with FAIMS for these abundance ranges, respectively. With No FAIMS the number was reduced to 1,10 and 61 for proteins and 2, 22, and 140 for peptides, respectively. This observation is pointing that FAIMS notably increase the detection of lower abundance proteins and peptide levels resulting in a vast dynamic range of ion detection. Therefore, coupling FAIMS with conventional nanoLC holds promise for enhancing the sensitivity and depth of proteomic profiling ^1^ ^41^ ^42^.

### Comparative analysis of SAP25 complex proteins with FAIMS and with No FAIMS configurations

Next, we sought to determine the abundance of Halo-SAP25 affinity-purified protein complexes, including the Sin3 complex and enzymatic protein complexes previously identified in our study^14^. Our focus here is to evaluate the abundance of these protein complexes both in the presence and absence of FAIMS. The enrichment of these proteins was determined by using normalized total abundance (log2) (Table S1). We compared these values for Sin3 complex proteins which includes: HDAC2, SAP25, BRMS1, ING2, SIN3B, ING1, SAP130, SIN3A, SAP30, RBBP7, HDAC1, BRMS1L, ARID4A, SUDS3, ARID4B and SAP30L (Figure 5A). We observed that ARID4A and BRMS1L were only observed with FAIMS, while ARID4B, SAP30L, and SUDS3 were present with No FAIMS configurations; however, this observation was limited to a single SAP25 replicate with abundance between 2.06E+05 to 6.09E+05 (Table S1). On the other hand, we did not observe major differences in the normalized total abundances (log2) with FAIMS and No FAIMS configurations for other SAP25 associated proteins. The only proteins having higher abundances were SKP1 and CUL1 while FBXO3, OGT and USP11 have higher abundances with the no FAIMS configurations (Figure 5B). This result suggests that FAIMS does not significantly affect the detection of SAP25 complex proteins due to their presence in higher abundances in samples compared to other proteins (Table S1).

**Figure 5.**
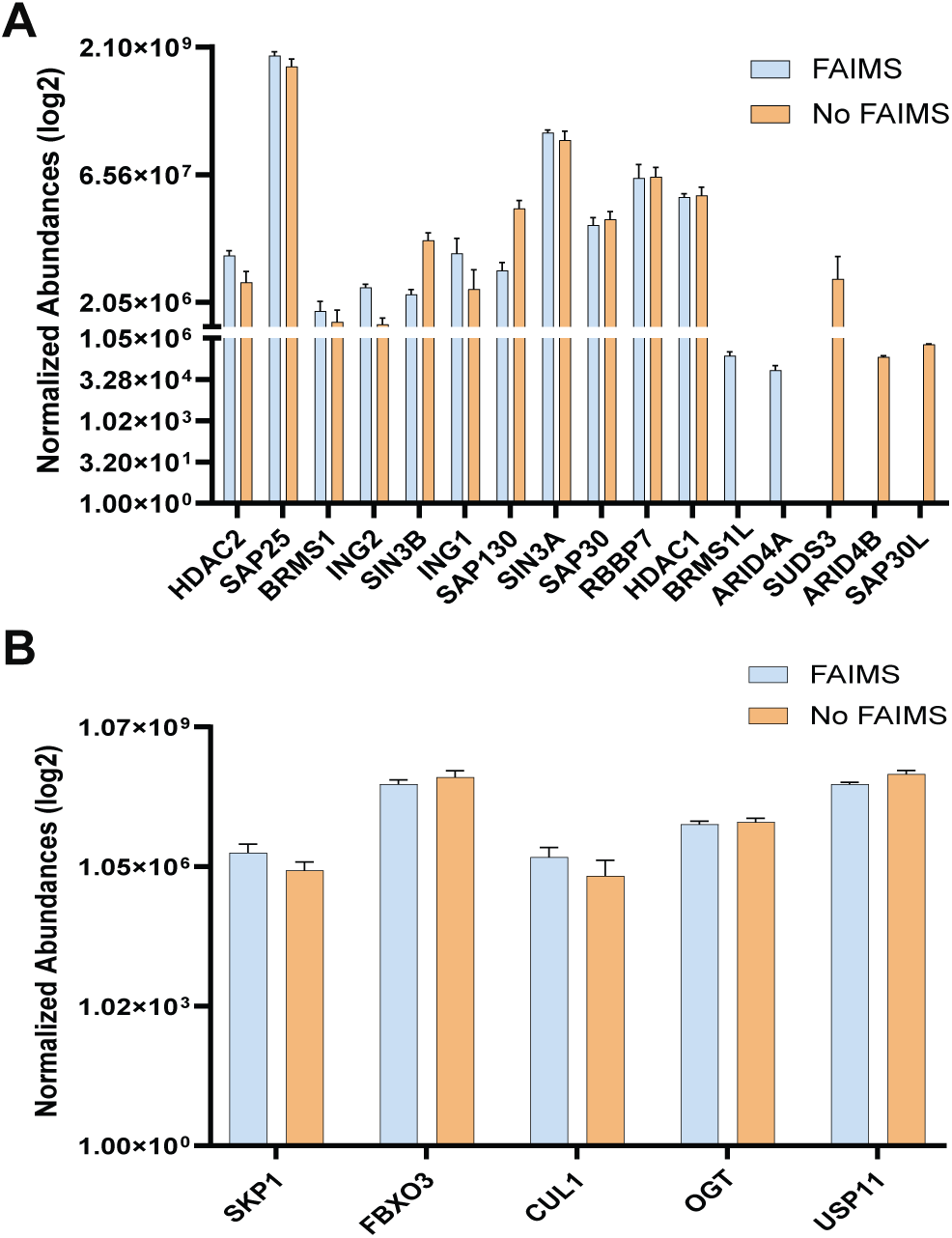
SAP25 binding partners under FAIMS and No FAIMS configurations. (A). Box & Whiskers plot showing Normalized protein abundances (log2) for Sin3 Complex proteins observed with SAP25 purification detected in FAIMS (Blue) and with No FAIMS (Orange) configurations. Values represent the averages of six runs. (B) Box & Whiskers plot showing Normalized protein abundances (log2) for Enzymatic and regulatory proteins observed with SAP25 purification detected in FAIMS (Blue) and with No FAIMS (Orange) configurations. Values represent the averages of six runs.

### FAIMS technology filtering potential contaminants in Affinity-Purified Protein Sample

Next, we evaluated the importance of FAIMS for detecting and filtering contaminants, commonly introduced during protein sample preparation using Skyline ^34^ ^35^ ^36^. We detected and quantified a total of 575 precursor ions to common chemical contaminants using Skyline, and we merged the multiple precursor ion abundances for most of the chemical contaminants into 23 total chemical contaminant abundances (Table S5-6). This included Tween and Triton series contaminants, and other detergents such as NP-40 and CHAPS, as well as Polyethylene Glycols (PEGs) and PEG Derivatives, Quaternary Ammonium Compounds and various other Organic and Polypropylene Glycols (PPG) and Derivatives (Table S5-6). By comparing the total contaminant abundances across twelve SAP25 runs (six for each configuration), we observed that most of the contaminants especially Tween series contaminants (Tween 20, Tween 40, Tween 60, and Tween 80), Triton X-100, IGEPAL-630, Didodecyl 3,3’ thiodipropionate and CHAPS showed substantial reduction in detection under FAIMS configuration.

For example, Figure 6 shows the chromatographic profiles of Triton X-100 without FAIMS (6A), Triton X-100 with FAIMS (6B), Tween 60 without FAIMS (6C), Tween 60 with FAIMS (6D), NP-40 without FAIMS (6E), and NP-40 with FAIMS (6F). Each of these chemical contaminants eluted late in the chromatography with Triton X-100 eluting starting at approximately 130 minutes (Fig. 6A-B), Tween60 eluting starting at approximately 116 minutes (Fig. 6C-D), and NP-40 eluting at approximately 114 minutes (Fig. 6E-F). For each chemical contaminant, the intensities were significantly higher in without FAIMS when compared to with FAIMS. The highest contaminant filtration was observed for Triton X-100, which is a reagent added to our high salt lysis buffer to lyse cells, where the level of filtration was significantly greater than 1000 times compared to the No FAIMS configuration (Fig. 6A-B). However, certain contaminants such as Dipalmityldimethylammonium chloride, Distearyldimethylammonium chloride, PEG, Phthalate esters, SPDMA and Triton X-100 reduced showing minimal filtering by FAIMS (Figure 7A and, Table S5-6).

**Figure 6.**
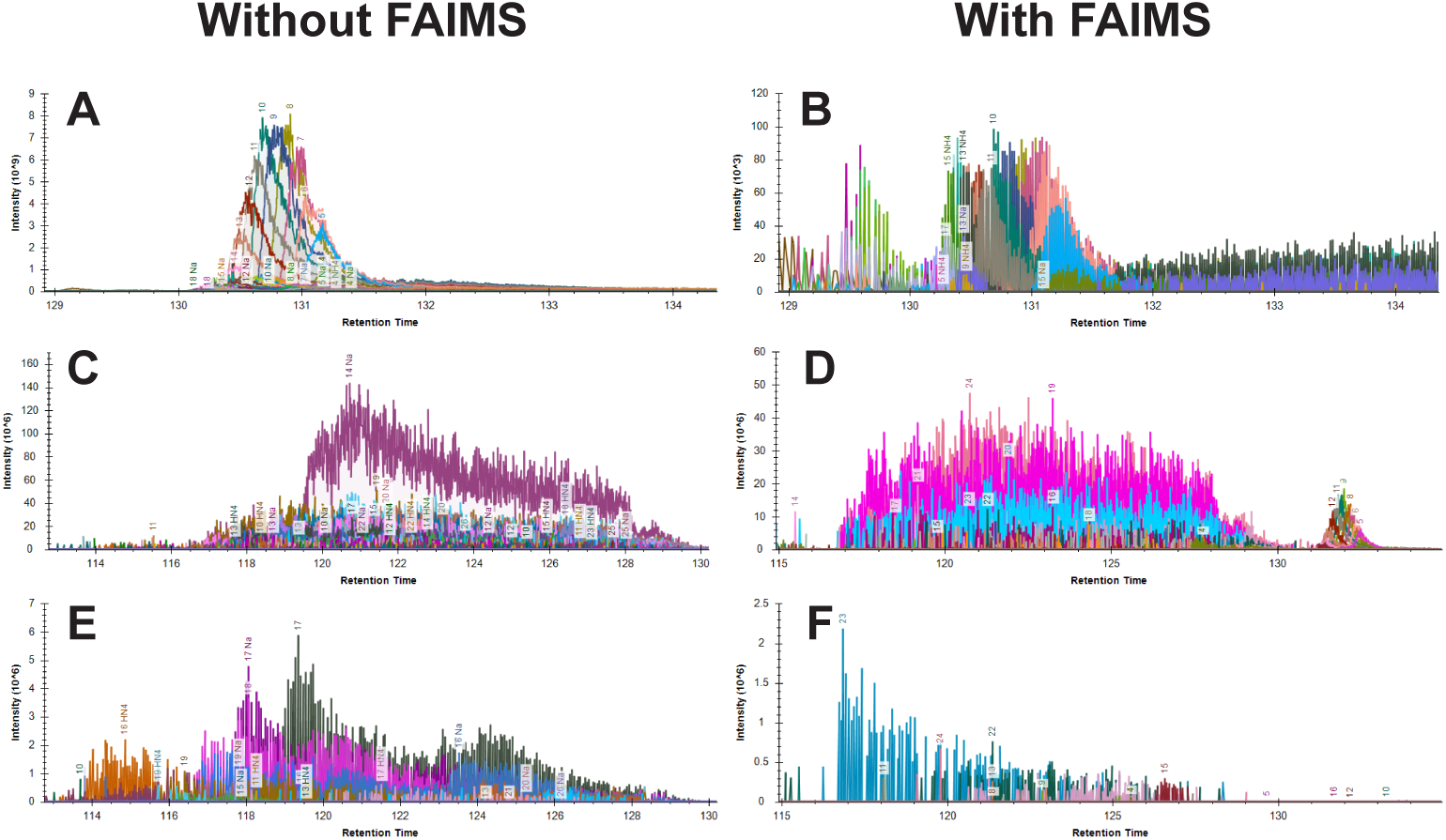
Chromatography of major contaminants within the LC-MS/MS analysis of SAP25 purifications. Extracted chromatograms visualized and analyzed in Skyline of several abundant precursor ions associated with Triton X-100 without FAIMS (A), Triton X-100 with FAIMS (B), Tween60 without FAIMS (C), Tween60 with FAIMS (D), NP-40 without FAIMS (E), and NP-40 with FAIMS (F). Within each pair of chromatograms for each molecule, different colors are associated with individual precursor ions detected.

**Figure 7.**
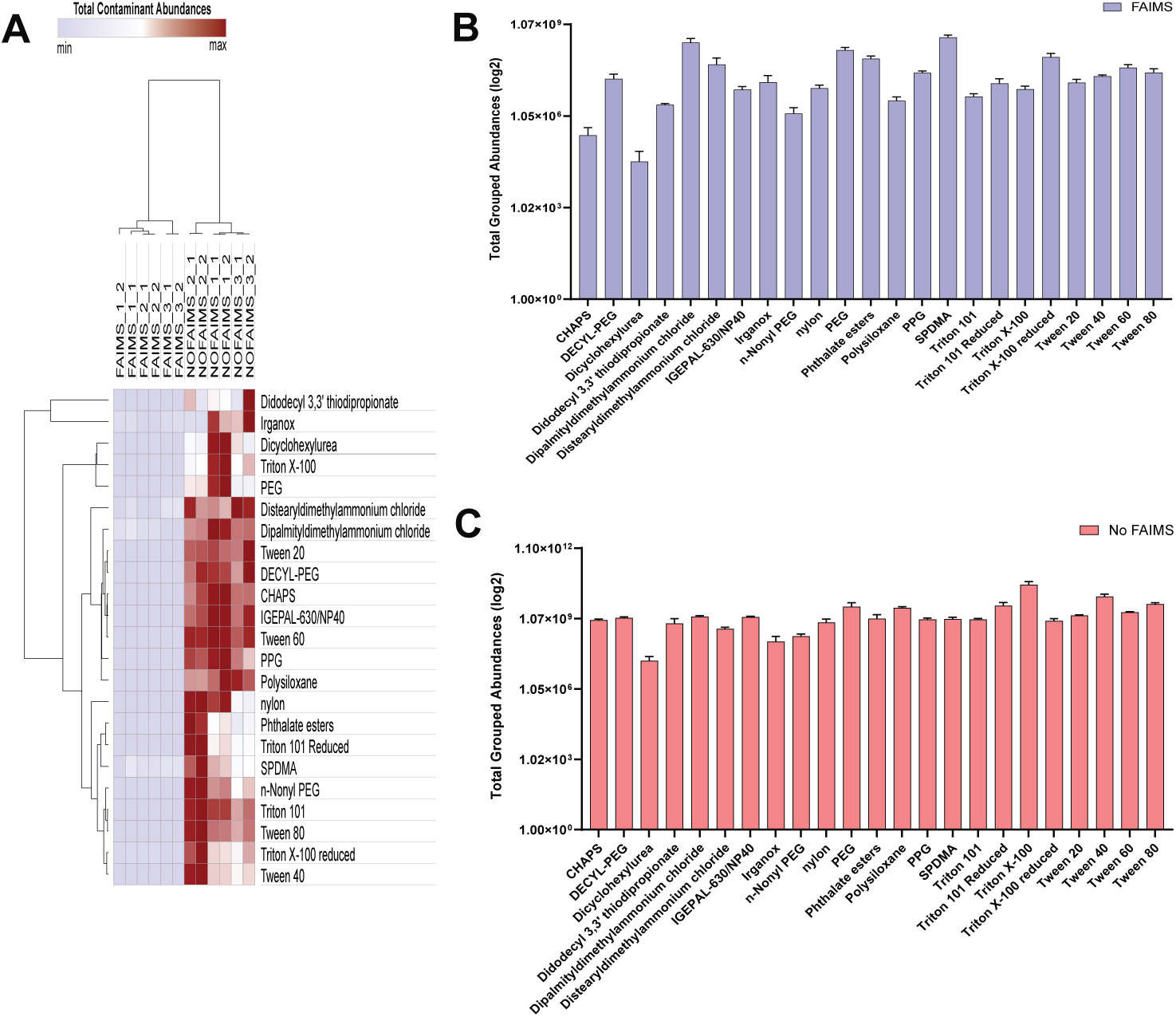
FAIMS significantly reduce the level of Contaminants. (A). Hierarchical clustering of Contaminants observed in Halo-SAP25 affinity purification with FAIMS and No FAIMS configuration. The comparative heatmap was created using the Morpheus online tool (https://software.broadinstitute.org/morpheus). Values were total contaminants abundance. Clustering was performed with One minus Pearson correlation and average linkage. (B). Bar chart of contaminants showing total grouped abundances observed with SAP25 FAIMS configuration. Values represent the averages of six runs. (C). Bar chart of contaminants showing total grouped abundances observed with SAP25 No FAIMS configuration. Values represent the averages of six runs.

Lastly, we decided to analyze the grouped abundances of contaminants (log2) with FAIMS and No FAIMS configuration. We used the average of contaminant abundances across six SAP25 runs for each configuration (Figure 7B-C). Our observation suggests that in the FAIMS configuration, a significant reduction in contaminant abundance is observed across various contaminants, many of them are having grouped abundance less than 1×10^7^ (Table S6). On the other hand, contaminants such as Dipalmityldimethylammonium chloride, PEG, SPDMA and Triton X-100 reduced having higher grouped abundances between 1×10^8^ to 5×10^8^ similar to No FAIMS configuration (Figure 7C), indicating the minimal effect of FAIMS to minimize these contaminants level. Overall, we determined that FAIMS played an important role in filtering various contaminants at significant levels, especially that of Triton X-100 (Figure 7B-C).

## Discussion

A significant challenge in proteomics experiments is the dynamic range of coeluting ions where some of these ions are not selected for fragmentation and are not peptides. In addition, ions with lower abundance can be outcompeted by higher abundance ions and this can potentially exceed instrument sensitivity. Many mass spectrometers select precursor ions with only high intensity and will often filter out low intensities of peptides ^43^. Therefore, methods and approaches to analyzed low abundance peptides from proteomics samples will enhance the data comprehensiveness and coverage.

FAIMS expands the dynamic range of LC-MS/MS analyses ^6, 23, 25, 44^. FAIMS increases signal-to-noise ratios by reducing the chemical background and acting as a separation device and generating cleaner mass spectra with higher confidence peptide detection ^25^. FAIMS readily separates multiply charged peptide precursors from singly charged background ions and can also separate based on charge state, conformers, and structural features ^17, 45^. Studies have shown that FAIMS enhances sensitivity when paired with monolithic capillary columns on an Orbitrap Fusion Lumos ^1^. Coupling the FAIMS interface with MS has shown significant promise in enhancing the sensitivity and depth of proteomic profiling ^1, 18, 26^, including in human serum ^23^, FFPE tissue, and exosomes ^6, 20^. Previously, there were several studies conducted with FAIMS on an Orbitrap Eclipse Tribrid mass spectrometer for Human Serum Proteoforms and single-cell proteomics ^23, 46^ and FLAG-tagged Co-IP analysis ^28^, Orbitrap Exploris 480 for FFPE Tissue and Plasma-Derived Exosomes ^6^ and, the Orbitrap Fusion Lumos mass spectrometer for targeted proteomics of tumor tissue ^47^. Beyond all these, the latest Orbitrap Ascend Tribrid Mass Spectrometer, which features a modified Orbitrap with additional ion-routing multipole (IRM) enhances scan speed and detection sensitivity, improves mass spectra quality, expands proteome coverage, and uncovers functionally important low-abundance proteins ^30, 31^. However, we were unable to find a study that compares the performance of the Orbitrap Ascend Tribrid Mass Spectrometer with FAIMS and without FAIMS for affinity-purified protein complexes.

Here we studied the benefits of FAIMS for the analysis of affinity-purified protein complexes, comparing the performance of Orbitrap Ascend Tribrid Mass Spectrometer with FAIMS and with No FAIMS. Identifying protein complexes and their subunits is a key to understanding many biological processes. Proteomic studies can identify a protein’s interaction partners, shedding light on the functions and regulatory mechanisms of protein complexes ^48^. This method allows for the exploration of protein interaction networks within a cell, offering critical insights into the structure and function of its protein machinery ^11^. Therefore, an in-depth protein analysis is crucial.

In this study, we found that mass spectrometry analyses with No FAIMS of affinity-purified SAP25 protein complex isolated from Flp-In™-293 cells resulted in fewer unique proteins and peptides, which is 13 and 169, respectively. Additionally, the average number of proteins and peptides detected among six SAP25 runs were less than 159 and 390, respectively. However, the number of proteins and peptides substantially increased when FAIMS was coupled to the Orbitrap Ascend Tribrid Mass Spectrometer, resulting in 92% and 77% gain in numbers for unique peptides and 53% and 43% for a total number of proteins and peptides respectively (Table S1-S2). The significant rise in the number of proteins and peptides for AP-MS samples with FAIMS is due to the ability of FAIMS to filter ions of interest at the entrance of the mass spectrometer ^6, 23, 26, 40^. In addition, the normalized total abundances for proteins and peptides observed with FAIMS and No FAIMS have a significant difference among the total six SAP25 runs. Despite that, when we consider the abundance of individual SAP25 runs, the increase in total normalized protein and peptide abundances at the log2 scale with No FAIMS was higher mostly due to the presence of high abundance proteins ^6, 21^. This suggests that using FAIMS with nanoLC-MS will strongly enhance the detection of proteins and peptides and is superior to traditional nanoLC-MS, as observed previously with U937 myeloid leukemia cells ^17^, HeLa cells^26^ and K562 cells ^18^.

One of the noteworthy results of the study was related to the effects of FAIMS on missing values, considered as a major issue in mass spectrometry data analysis ^37^ ^38^ ^39^ by hindering the statistics and reproducibility of data ^49^. Previously it was determined that using FAIMS with mass spectrometry significantly reduces the number of missing values ^6, 40^. The same observation occurs with our AP-MS study. We have observed that FAIMS significantly reduced the number of missing values compared to the No FAIMS configuration, therefore, significantly increasing the number of peptides and proteins identified and quantified (Figure 4A,4B, and 4C). The missing values in No FAIMS group were mostly related to low abundances at the peptide and protein levels. Importantly, the proteins and peptides with low abundances were identified and quantified exclusively under FAIMS configurations (Figure 4D, 4E), which may be lost by conventional LC-MS analyses. This observation suggests that FAIMS significantly improves signal-to-noise (S/N) ratio and filtering ions based on distinct mobility resulted in the identification of low abundance proteins and peptides ^25, 44^.

Finally, a potential limitation of the study is the work was carried out with three CVs and the default FAIMS configuration. We found that FAIMS with three CVs with DDA acquisition provides the highest protein and peptide-level detection. Further, we determined that FAIMS minimizes missing values and filtered the contaminants ^37^. However, some SAP25-associated proteins (Figure 5) detected under the No FAIMS configuration were lost with FAIMS. This may be attributed to FAIMS self-cleaning mechanism, field heating effects or limited ion transmission ^25, 50^. Therefore, further experiments may be required to optimize FAIMS parameters and the right selection of CVs value, which could lead to a better outcome by enhancing detection of proteins and peptides ^17, 24, 45^. Furthermore, optimizing both chromatography and mass spectrometry parameters may enhance the separation and detection at protein and peptide levels.

## Conclusion

The FAIMS source, when coupled between nanoLC and Orbitrap Ascend Tribrid Mass Spectrometer (nanoLC-FAIMS-MS) improves the dynamic for bottom-up proteomics, including affinity-purified protein complexes. Here, we demonstrated that the application of FAIMS with three CVs on an Orbitrap Ascend Tribrid Mass Spectrometer in DDA increased the sensitivity for protein and peptide detection by 42% and 44% respectively compared to No FAIMS configuration. More specifically, FAIMS enhances the sensitivity of nanoLC by reducing the level of chemical contaminants and reducing the number of missing values from 60.17% (No FAIMS) to 14.57% (FAIMS). Therefore, our study suggests FAIMS is a valuable component to proteomics analyses. Overall, our result indicates that FAIMS addresses several limitations of AP-MS based proteomics analysis at chemical contaminant and peptide levels. One potential approach for researchers to use when they do not have FAIMS on their system would be to use sample preparation approaches or digested sample clean up approaches that would reduce the chemical contaminants of their sample of interest prior to loading it onto an LC-MS/MS system.

---

**Figure S1.**
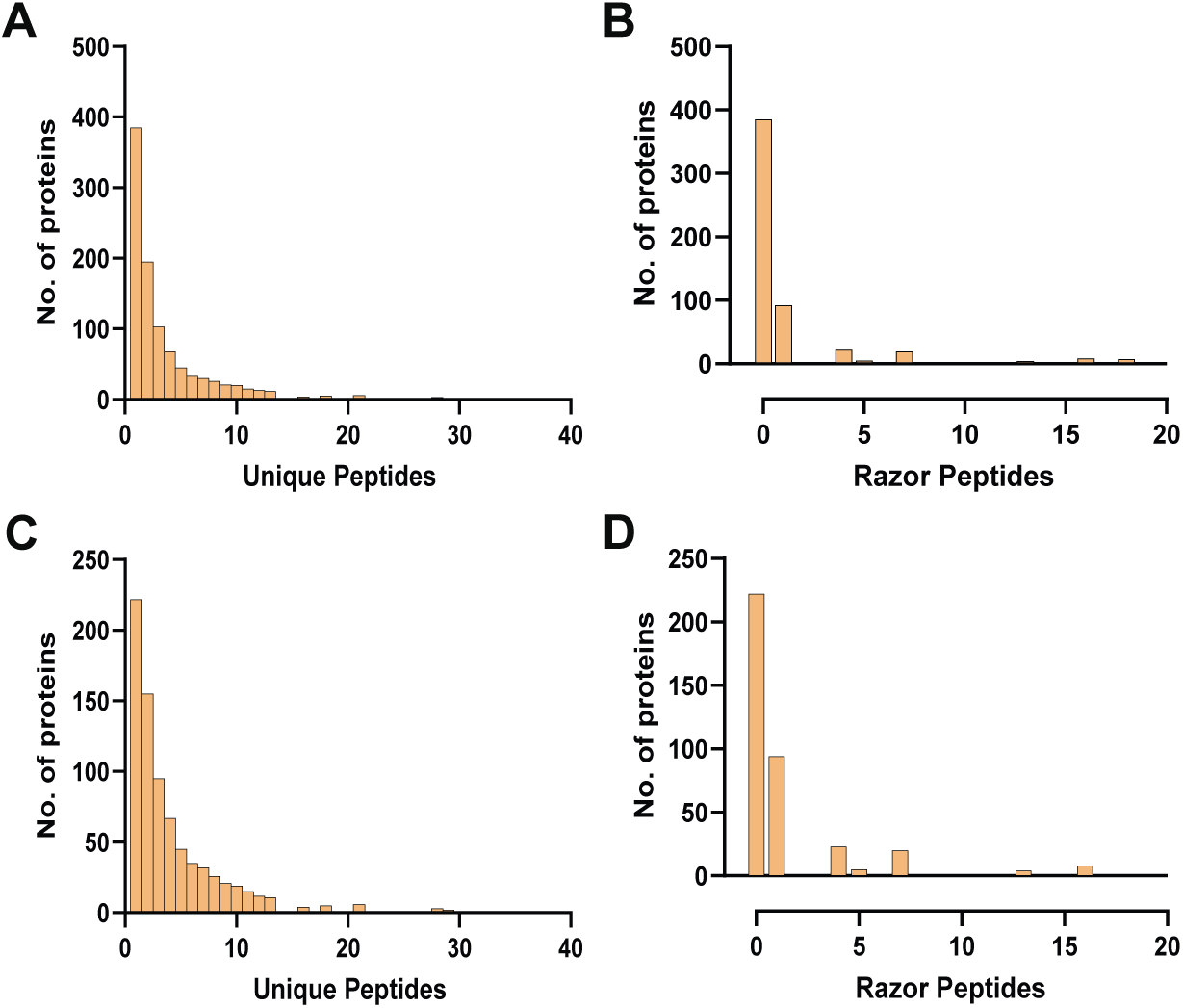
Distribution of razor and unique peptides under FAIMS and No FAIMS configurations. (A) Distribution of unique peptides with FAIMS. (B) Distribution of razor peptides with FAIMS. (C) Distribution of unique peptides with No FAIMS. (D) Distribution of razor peptides with No FAIMS.

## Supporting Information

The supporting data tables file includes:

Detailed information for Protein level detection observed with SAP25 Affinity purification (Table S1), Detailed information for Peptide level detection observed with SAP25 Affinity purification (Table S2), Detailed information about log2 value for Protein level detection (Table S3), Detailed information about Unique and razor peptides observed with SAP25 Affinity purification (Table S4), Detailed information about the Contaminants derived from Skyline (Table S5), and Detailed information about the Contaminants observed with SAP25 Affinity purification (Table S6).

## Data availability

The mass spectrometry data form the 6 FAIMS and 6 No FAIMS runs have been deposited in the MassIVE repository (http://massive.ucsd.edu) with the data identifier MSV000096548 (ftp://massive.ucsd.edu/v07/MSV000096548/).

## Author Contributions

Conceptualization: PG, MR, MPW

Investigation: PG, JC, MR, ZC

Formal Analysis: PG, JC, MR, ZC

Resources: PG, JT

Data Curation: PG, JC, MR, ZC

Writing: PG, JC, MPW

Supervision: MPW

Funding acquisition: MPW

## Notes

The authors declare that they have no conflicts of interest related to the work presented here.

## Acknowledgements

The research presented here was supported by the National Institute of General Medical Sciences of the National Institutes of Health R35GM145240 (M.P.W.). The content is solely the responsibility of the authors and does not necessarily represent the official views of the National Institutes of Health. In addition, J.C. receives support from the Stowers Institute for Medical Research.

## For Table of Contents Only

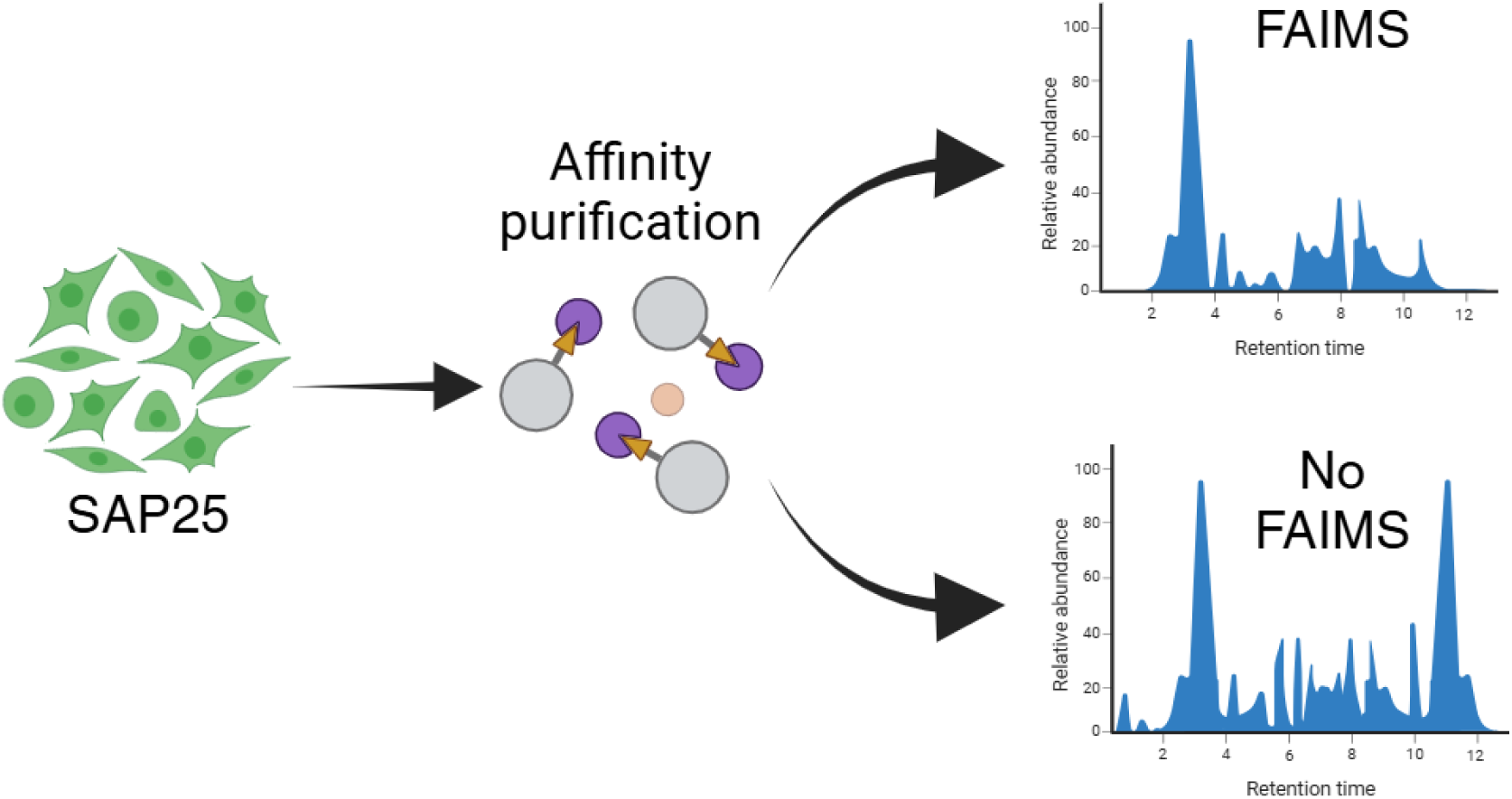

